# FastQTLmapping: an ultra-fast package for mQTL-like analysis

**DOI:** 10.1101/2021.11.16.468610

**Authors:** Xingjian Gao, Jiarui Li, Xinxuan Liu, Qianqian Peng, Han Jing, Sijia Wang, Fan Liu

## Abstract

**Summary:** Here we describe fastQTLmapping, a C++ package that is computationally efficient not only for mQTL-like analysis but as a generic solver also for conducting exhaustive linear regressions involving extraordinarily large numbers of dependent and explanatory variables allowing for covariates. Compared to the state-of-the-art MatrixEQTL, fastQTLmapping was an order of magnitude faster with much lower peak memory usage. In a large dataset consisting of 3,500 individuals, 8 million SNPs, 0.8 million CpGs and 20 covariates, fastQTLmapping completed the mQTL analysis in 7 hours with 230GB peak memory usage.

**Availability and Implementation:** FastQTLmapping is implemented in C++ and released under a GPL license. FastQTLmapping can be downloaded from https://github.com/TianTTL/xQTLmapping.

## INTRODUCTION

Methylation quantitative trait loci (mQTL) analysis is to test association between a large number of genomic variants and a large number of CpG sites over the genome, ideally using well-sized population samples to obtain sufficient statistical power, with covariates to control for potential confounding effects, and in an exhaustive manner to maximize genome resolution. Such analysis is often highly computationally burdensome, easily involving trillions of multiple regressions. MatrixEQTL^1^ represents the state-of-the-art in terms of computational efficiency, yet has room for improvement.

Here we describe fastQTLmapping, a computationally efficient, exact, and generic solver for exhaustive multiple regression analysis involving extraordinarily large numbers of dependent and explanatory variables with covariates, which is particularly helpful in mQTL-like analysis.

## IMPLEMENTATION

FastQTLmapping accepts input files in text format and in Plink binary format. The output file is in text format and contains all test statistics for all regressions, with the ability to control the volume of the output at preset significance thresholds. Different thresholds can be specified according to physical distances between the markers under investigation, which facilitates the analysis of cis- and trans-mQTLs. Z-and rank-normalizations are optional for pre-processing certain or all input variables. In order to maximize variable retention, fastQTLmapping determines missing values when concatenating individual dependent and explanatory variables, which in turn is quality controlled at a user-specified threshold. FastQTLmapping is deployed on Linux using MKL (https://software.intel.com/tools/onemkl) and GSL (http://www.gnu.org/software/gsl/) library, and is run from the command line. C++ source code, an example run, and documentation are freely available at https://github.com/TianTTL/xQTLmapping.

FastQTLmapping loads the entire input file into the memory as characters and then converts them into floating point numbers. A floating-point parser is developed to effectively handle different data types in parallel on a per-line basis.

Redundant calculations related to covariates are removed using an orthogonal analysis. Consider a multiple regression, ***C*** is the matrix of *k* covariates,

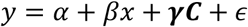

orthogonalizing *x* and *y* with respect to ***C***. Decompose ***C*** using QR factorization ***C*** = ***QR***, then centralize ***Q*** to 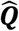, where 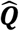 is an orthonormal basis. Then subtract the projections of *x* and *y* on 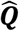 to get *x*′ and *y*′.

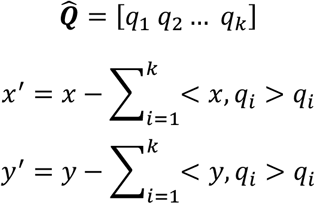

The subsequent analyses only involve univariate regressions *y*′ = *α*′ + *β*′*x*′ + *ϵ*′ to test *H*_0_: *β*′ = 0 with *k* less degrees of freedom. FastQTLmapping speeds up the pair-wise univariate regressions by calculating Pearson product-moment correlations, obtains the p-values based on the t-distribution, and produces other test statistics if a p-value satisfies a preset significant threshold.

Mathematical operations are accelerated using the Math Kernel Library (MKL), serial computations are parallelized using OpenMP, and peak memory consumption is controlled through data splitting and variable reuse. Omics data are divided into blocks and dynamically assigned to threads. The block sizes are self-adapted based on the data size to maximize the performance of Level-3 BLAS while controlling for memory consumption. Critical section is used to guard smooth writing from threads to hard disks. All intermediate variables are allocated as aligned memory buffer.

## PERFORMANCE

We compared the performance of fastQTLmapping and MatrixEQTL. For a fair comparison, we made a parallel version of MatrixEQTL using R packages ‘doParallel’, linked MKL to R environment, and manually split data to feed MatrixEQTL to achieve its optimal performance. The test data was downloaded from the GEO database (https://www.ncbi.nlm.nih.gov/geo/). The SNP and CpG data are GSE79254 and GSE79144, respectively as described previously.^2^ The original data contains 54 individuals, 1.5×10^6^ SNPs, and 4.5×10^5^ CpGs. For testing purpose, we individual-wise resampled the data with replacement and generated 4 datasets consisting of 100, 200, 300 and 400 individuals. MQTL analysis was conducted with single CPU thread and with 8 CPU threads without covariates. The testing environment is CPU: Xeon E5 2686 V4 (18 cores, 2.3GHz), RAM: 256 GB ECC DDR4, OS: CentOS Linux 7, g++ version: 4.8.5, R version: 4.0.3, and MKL version: 2021.3.0. Both fastQTLmapping and MatrixEQTL produced the same and exact results in the absence of missing values under all investigated settings. In presence of missing values, both packages provided approximate results, while fastQTLmapping results were always closer to the exact results. The operating time of both fastQTLmapping and MatrixEQTL was largely linear to the data size. For computation, fastQTLmapping was 9.0-19.2 times faster than MatrixEQTL under the single-thread setting and 9.2-16.9 times faster under the 8-threads setting (**Figure 1A**). For I/O, fastQTLmapping was 19.6-34.7 times faster than MatrixEQTL under the single-thread setting and 11.6-16.8 times faster under the 8-threads setting (**Figure 1B**). The peak memory consumption of fastQTLmapping (1.7-4.9 GB) was much smaller than that of MatrixEQTL (9.7–13.7 GB) under the single-thread setting. The increase of the peak memory consumption when parallelizing to 8 threads was slower for fastQTLmapping (1.5-2.5 folds) than MatrixEQTL (5.7-6.9 folds, **Figure 1C**).

**Figure 1.**
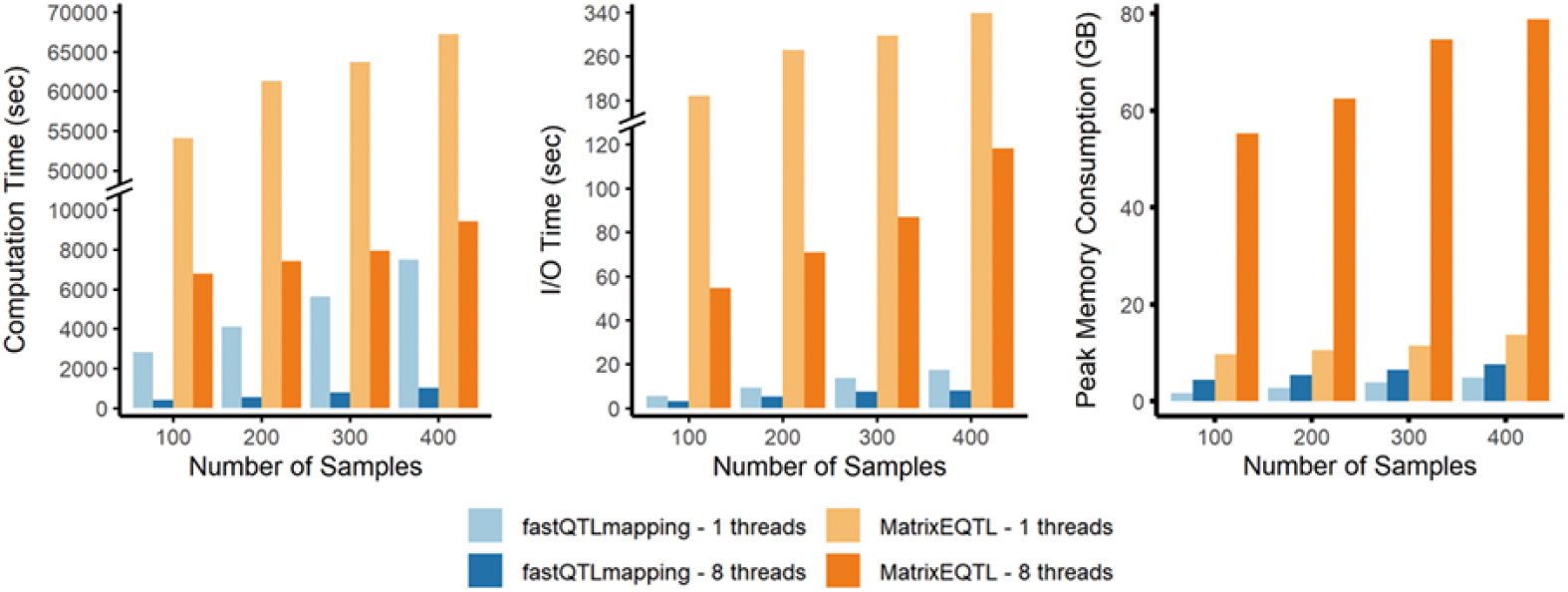
Performance of fastQTLmapping and MatrixEQTL under various settings. A, Computation time; B, I/O time; C. Peak memory consumption.

To further examine the performance of fastQTLmapping in well-sized population studies using densely imputed SNPs and Illumina MethylationEPIC 850K BeadChip, we also generated a dataset consisting of 3,500 individuals, 8×10^6^ SNPs, and 8×10^5^ CpGs by individual-wise, SNP-wise, and CpG-wise resampling the original data with replacement, and added 20 normally distributed variables as covariates. With 20 threads running in parallel, fastQTLmapping completed the mQTL analysis including I/O in 7.0 hours with 230 GB peak memory.

## CONCLUSIONS

FastQTLmapping is ultra-fast, easy-to-deploy, capable for conducting pair-wise regression analysis at extraordinarily large scales on regular servers, particularly helpful for well-sized mQTL studies.

## Acknowledgements

This work was supported by the Strategic Priority Research Program of Chinese Academy of Sciences [Grant No. XDB38010400, XDB38010100, XDC01000000], Shanghai Municipal Science and Technology Major Project [Grant No. 2017SHZDZX01], National Natural Science Foundation of China (NSFC) [81930056, 91631307], Science and Technology Service Network Initiative of Chinese Academy of Sciences [KFJ-STS-QYZD-2021-08-001, KFJ-STS-ZDTP-079], the National Key Research and Development Project [Grant No. 2018YFC0910403], the Max Planck-CAS Paul Gerson Unna Independent Research Group Leadership Award, and the CAS Youth Innovation Promotion Association [Grant No. 2020276].

